# Graph regularized, semi-supervised learning improves annotation of *de novo* transcriptomes

**DOI:** 10.1101/089417

**Authors:** Laraib I. Malik, Shravya Thatipally, Nikhil Junneti, Rob Patro

## Abstract

We present a new method, GRASS, for improving an initial annotation of *de novo* transcriptomes. GRASS makes the shared-sequence relationships between assembled contigs explicit in the form of a graph, and applies an algorithm that performs label propagation to transfer annotations between related contigs and modifies the graph topology iteratively. We demonstrate that GRASS increases the completeness and accuracy of the initial annotation, allows for improved differential analysis, and is very efficient, typically taking 10s of minutes.

---

Advances in sequencing technologies have allowed the efficient and accurate exploration of transcriptomes beyond the scope of genetic model organisms [1, 2]. The first major step in pipelines involving non-model organisms is often *de novo* transcriptome assembly. *De novo* assembly allows for the identification of novel transcripts, as well as transcript quantification and, in turn, differential expression studies across various conditions and cell types [3, 4]. However, for meaningful interpretation of these analyses, we must have some notion of what the contigs in our assembly represent. Often, we have annotated genomes from species related to these non-model organisms. This information can be harvested to accurately annotate the contigs of a *de novo* assembly and improve our understanding of down-stream analyses. Traditionally, variants of BLAST [5] are used to perform this annotation and then complete Gene Ontology (GO) analysis [6, 7]. Apart from these, other methods are used for pre-processing the assembly in order to filter out spurious contigs [8]. However, a large proportion of contigs may remain unannotated even after these steps.

We present a method, GRASS (Graph Regularized Annotation via Semi-Supervised learning), that employs data from previously annotated species and transfers transcript and gene labels to the current assembly (overview in Fig.1). This, in turn, improves annotation quality. The annotation performed by GRASS is done in conjunction with a graph that represents contig-level similarity in the *de novo* assembly. In this sense, GRASS is different from existing annotation methodologies, since it takes advantage not only of similarity between the transcripts of related species, but also considers sequence similarity between contigs *within* the *de novo* assembly. We consider a weighted graph, *G*, in which the contigs are the vertices and each pair of contigs is connected by an edge if reads multimap between them. The weight of each edge is proportional to the fraction of multimapping fragments between the adjacent contigs. This graph can be efficiently built using previously generated fragment equivalence classes (which we derived from the output of Sailfish [9, 10] or Salmon [11]). These classes represent the relationship between fragments based on the set of transcripts to which they map or align. This concept has proven very powerful in transcript-level quantification, and was first introduced, to the best of our knowledge, by Turro et al. [12]. The efficient construction of this graph is implemented in RapClust [13], a tool for clustering contigs in *de novo* transcriptome assemblies.

**Figure 1:**
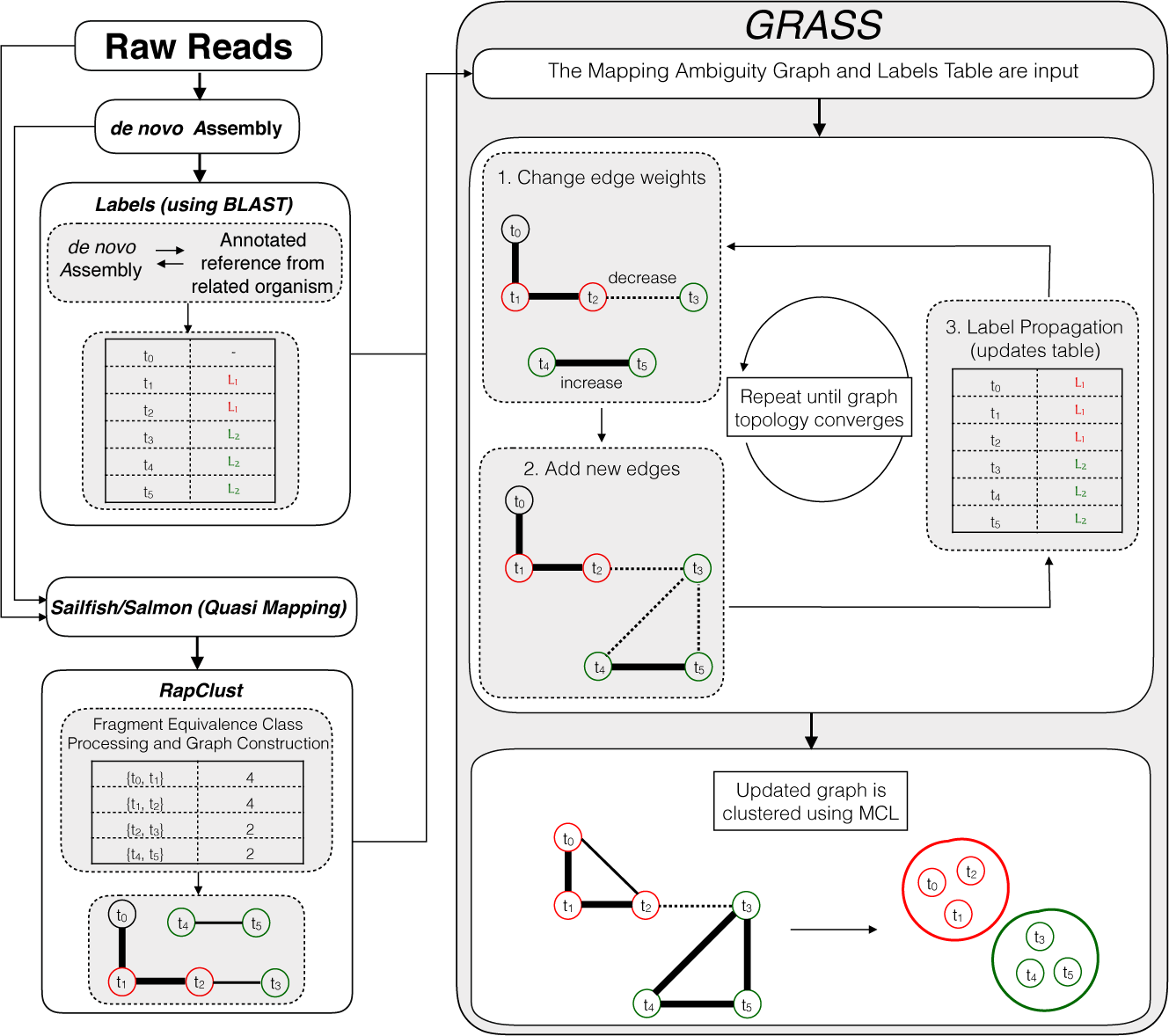
GRASS requires two inputs. The first is a partial contig to label mapping (i.e., a labeling of some subset of contigs with gene names), which can be obtained using an existing method such as performing a BLAST search of the contigs against a closely related species. The second input is a mapping ambiguity graph constructed using the previously generated equivalence classes [13]. The first step of the iterative algorithm in GRASS modifies the edge weights and adds new edges based on the current labeling (linking contigs that share a gene label with high probability), and the second step updates the labels using a label propagation method [14]. These steps are repeated until the number of edges in the graph converges.

We note that, rather than being a complete annotation pipeline, GRASS is best considered as a method capable of “boosting” an initial set of annotations by accounting for expression and sequence similarities within the *de novo* assembly. Furthermore, GRASS not only boosts annotation quality, but uses these annotations to improve contig-level clustering, which can lead to more accurate differential analysis results (see Figure 2). As such, in GRASS, we begin by labeling nodes of this graph, *G*, using a traditional approach, such as a BLAST search. Subsequently, a graph based semi-supervised learning method for label propagation is used to transfer these initial annotations to unannotated nodes in the graph. For this purpose, we use the adsorption algorithm, which relies on random walks through the graph and has been used to efficiently propagate information through a variety of graphs in its various applications [14]. On top of this label propagation, we build an iterative algorithm to modify the topology and edge weights in the graph based on the current labeling. This process is repeated until the topology of the graph converges. The final result of our approach is a collection of annotations for the contigs in the *de novo* assembly and a graph that best represents the relationship between these contigs based on the available sequence and annotation information (see Online Methods for details).

**Figure 2:**
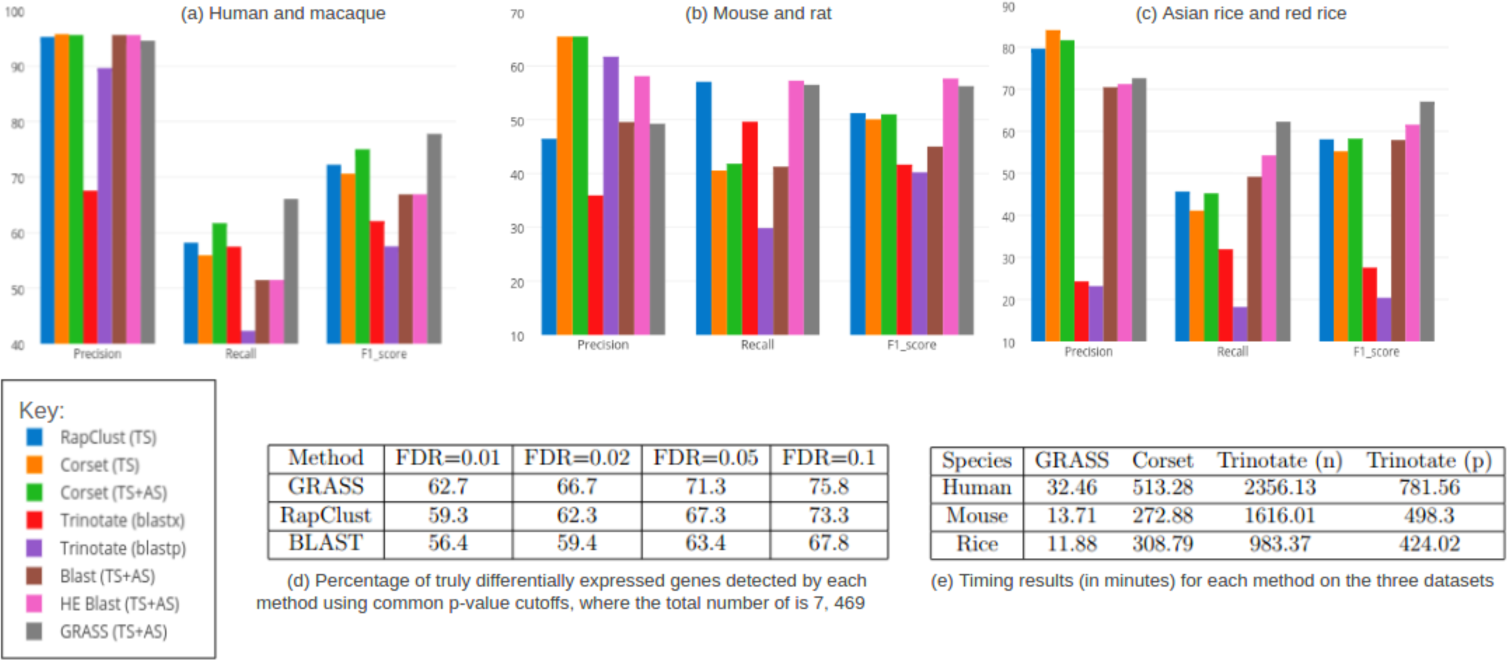
(a,b,c) Precision, recall and F1 scores for the three test species (TS) using a closely related annotated species (AS). Explanation for the key: 1. RapClust run using the processed output from Sailfish on the *de novo* assembly of the test species and RNA-seq reads. 2. Corset run using the *de novo* assembly of the test species and aligned reads. 3. Corset run using the *de novo* assembly of the test species and aligned RNA-seq reads to both the assembly and annotated genome of the related species. 4. Nucleotide BLAST between contigs from the *de novo* assembly of the test species and the SwissProt database. 5. Protein BLAST between contigs from the *de novo* assembly and the SwissProt database. 6. Nucleotide BLAST between all the contigs in the *de novo* assembly of the test species and the annotated genome from the related species. 7. Nucleotide BLAST between just the expressed contigs in the *de novo* assembly and the annotated genome from the related species. (Contigs with <10 reads mapping to them are removed.) 8. GRASS using the output from Sailfish and contigs in the *de novo* assembly labeled using the annotated genome from a related species. (d) GRASS is several times faster than the other tools. (e) GRASS improves the rate of discovery of differentially expressed genes compared to th other methods.

We compare our results against a number of other approaches, some of which are used for integrating information from annotated species with *de novo* assemblies. Since there is no obvious and complete mapping between the contigs and annotations from the related species, we choose to compare the different methods based on their ability to give contigs from a single gene the same label and, in turn, cluster them together for down-stream analyses. The first is RapClust [13], which does not make use of any annotations to cluster contigs in the *de novo* assembly. We test Corset [15] in two ways, with and without the annotated related species. We also show a comparison against clustering that is based on the label assigned to each contig after a simple nucleotide BLAST of the *de novo* assembly against the reference transcriptome. This is done for both the complete set of contigs, as well as only the contigs that have more than 10 reads mapping to them (a criterion used to filter contigs in Corset, RapClust and GRASS). Note that this information is obtained from the quantification results of Sailfish, and is automatically incorporated in the mapping ambiguity graph of GRASS. The last set of results is obtained in a similar way (only on contigs with 10 or more mapping reads) after a BLAST search (both nucleotide and protein) against the release of SwissProt database used by Trinotate, but with the specific test species annotations removed [16] (http://trinotate.github.io/). For all analyses, reference-based clustering (obtained by BLASTing the contigs against the true reference genome) is treated as the truth, and GRASS is run using an *α* value of 0.8 (Supplementary Fig.1). We test on three main datasets, and show that GRASS provides a better clustering and labels a greater number of contigs, while also being substantially faster.

We assess the quality of our clusters based on precision and recall scores, where a pair of contigs is considered a true positive when they share a gene label under the true labeling, obtained using the annotated reference from the specific species. Co-clustered contigs sharing different ground-truth labels are considered as false positives, and identically-labeled contigs that reside in different clusters are considered as false negatives (all remaining pairs are true negatives). We show that our method performs much better than the others in terms of recall, while suffering only a minor loss compared to the next-best method in terms of precision (Fig.2(a,b,c)) (more results in Supplementary Fig.2 and 3). Integrating quantification results with BLAST and removing low expression contigs from the clustering improves the accuracy of BLAST-based clustering and annotation in some cases (Fig.2(b)). However, the extra quantification information is obtained by running Sailfish on the input RNA-seq data, and is automatically integrated into the methodology of GRASS. Improved clustering, in turn, also results in improved differential expression analysis. We verify this by demonstrating that GRASS, which iteratively integrates contig-level similarity and label annotations to improve contig clustering, outperforms RapClust, which clusters contigs using just the fragment ambiguity graph, and a BLAST-based clustering which uses only the annotations alone to define clusters. Specifically, we examine the rate at which differential expression calls made under the GRASS, RapClust and BLAST clusterings recover genes called as DE under the ground truth labeling (Fig.2(d)). We show that GRASS obtains a higher sensitivity at a given FDR than RapClust, and that both of these approaches achieve a considerably higher sensitivity at a fixed FDR than BLAST (Supplementary Fig.4). With an FDR of 0.05, GRASS is shown to have a higher recall rate than the other methods (Fig.2(d)).

In addition to being accurate, GRASS assigns annotations to a greater number of contigs than the other methods. This is a benefit of the label propagation step which, in each iteration, makes use of the previous information to continue labeling connected components in the graph. This means that some of the previously unannotated contigs, which could not be annotated on the basis of sequence similarity alone, are labeled by the end of the iterative process. In this way, GRASS is able to assign annotations to 1, 300, 1, 121 and 2, 177 extra contigs in human, mouse and rice, respectively. To assess the quality of this annotation, we check how these contigs co-cluster according to the newly-assigned labels, compared to their true annotation (Supplementary Table. 1) The larger number of annotations could be helpful in down-stream analyses for specific genes.

Along with producing more complete and accurate annotations, GRASS is also much faster than the other tools (Fig.2(e)). The time taken for GRASS includes the time to run Sailfish (which generates contig-level abundance estimates), to generate fragment equivalence classes, to construct the mapping ambiguity graph (using RapClust), to run BLAST to generate the initial contig labels, and then to propagate the labels and alter the graph topology using the GRASS algorithm itself. Similarly, Corset timing includes time to align reads in each sample against the reference using Bowtie [17]. GRASS takes a little over half an hour in human (for details of the experimental setup, refer Experimental Setup in online methods), with a total of 23.2 million reads across 6 samples and 107, 389 contigs in the *de novo* assembly. The other species take less than 15 minutes, with a total of 10.5 million reads and 75, 727 contigs in mouse and 8 million reads and 99, 745 contigs in rice. In comparison, the other tools take several hours to run on these datasets.

GRASS is able to combine information from three main sources: contig sequence similarity in the *de novo* assembly, quantification results after read mapping (used to remove spurious and very-low abundance contigs), and annotations from a closely related species. It does so much faster than other tools, and can produce substantially more accurate annotations, especially as the phylogenetic distance between the assembly and the reference species used for annotation grows. GRASS simplifies the process of annotating contigs from *de novo* assemblies, improves contig-level clustering, and introduces what we hope will be a powerful new idea in improving annotation quality and completeness.

## Software Availability

GRASS is written in Python, and is freely-available under an open-source (BSD) license at https://github.com/COMBINE-lab/GRASS.

## Acknowledgements

We gratefully acknowledge support from NSF grant BBSRC-NSF/BIO-1564917.

## Contributions

L.I.M. and R.P. designed the GRASS algorithm, and devised the experiments performed in the manuscript. L.I.M. implemented the algorithm. L.I.M., S.T. and N.J. conducted the experiments and prepared the figures and results. L.I.M. and R.P. wrote the manuscript.

## Online Methods

### Algorithm

GRASS is an algorithm for improving and annotating *de novo* assemblies using data available from related species that have already been annotated. These annotations are initially mapped to contigs in the *de novo* assembly using a pre-existing method and are passed to GRASS as input. We use BLAST for this purpose in all our tests, but other procedures may be used for the initial labeling. The other main input to GRASS is the mapping ambiguity graph constructed on the basis of fragment equivalence classes (we obtain this via RapClust [13]). An equivalence class is defined as a collection of reads, all of which map to exactly the same set of contigs. RapClust constructs a weighted graph using these classes where the vertices represent the contigs and a pair of contigs is connected if they co-label an equivalence class. This graph is then updated in GRASS and annotation information propagated through it.

The initial labeling can be passed to GRASS in two formats. Take A to be the set of contigs from the *de novo* assembly, and B the set of annotated transcripts from the related species. The first is a simple tab-delimited, two-column format with the contigs from the assembly in the first column and their respective labels from the related species in the second; essentially a mapping from A to B. The second format consists of two separate files where the first file contains results from a nucleotide BLAST of the *de novo* assembly of the test species against the database constructed using the annotated reference from the related species and vice-versa in the second file. Hence, the first file contains a mapping from A to B and the second from B to A. In the latter case, the two BLAST files are sorted using the bit score, with ties being broken by the e-value, and contigs are given their corresponding consensus label. The contig is not labeled if no consensus exists, i.e., if the best hit in A→B is not the same as the best hit in B→A. This labeling and the mapping ambiguity graph are passed to GRASS. Then, GRASS proceeds executing, iteratively, the steps of its algorithm include modifying the edge weights in the graph based on the annotation labels of the contigs and running a learning algorithm on the graph to annotate unlabeled contigs. These are explained in more detail below.

- G*^t^* = (*V, E^t^*) is the mapping ambiguity graph at the *t*^th^ iteration of the algorithm
- E*^t^* is the edge set at the t^th^ iteration
- *e* ∈ *2* E*^t^* is an edge from E*^t^* and is an un-ordered pair {*u*, *v*} | *u, v ∈ V*

#### Edge manipulation

After labeling contigs, edges in the graph are changed, iteratively, in two ways based on the shared labels and edge weights from prior iterations of the algorithm:

1. Let t + 1 denote the current iteration of the algorithm, and let *e* = {*u, v*} ∈ *E^t^* be an edge in the set *E^t^* of edges. The weight of *e*, denoted as *w* (*e*), can be updated if there are labels common between two contigs sharing an edge and is calculated as follows:

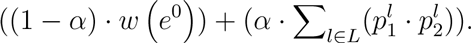

Here, *α* is set to 0.8 for our tests, *w*(*e*^0^) is the original edge weight in the input graph (i.e., *G*^0^) and *p*_1_, *p*_2_ are probabilities of each contig having a label l from the set of shared labels, *L*.
2. New edges are added to the graph in cases where two contigs share a label with high probability, but do not have an edge between them. The new edge weight is calculated in the same way as above. However, instead of the original edge weight *w* (*e*^0^) (since no edge originally existed), we instead use the median of the edge weights connecting the two vertices to their neighbors in the graph. The edge is only added if the joint probability of shared labels is greater than 0.9. This threshold is chosen to avoid adding a large number of false edges, especially in the first iteration when a majority of labels have an assigned probability of 1.0 (since this is the seed probability with which these labels are initialized).

#### Label propagation

Using results from BLAST, a portion of the contigs in the mapping ambiguity graph are labeled. The graph-based semi-supervised learning algorithm, *absorption* [14] (we use the implementation from the Junto library [20]), is used to extend these labels to contigs that have a large number of overlapping mapped reads and, therefore, an edge between them in the graph. The algorithm works by taking random walks through the graph carrying label information. This information is propagated based on three probabilities associated with each node: *p^inj^*, the probability of stopping the current random walk and emitting a new label, *p^abnd^*, the probability of completely abandoning the walk, and *p^cont^*, the probability of continuing the random walk with the current label.

We do not run the algorithm until convergence, since we wish to utilize information at each iteration to update the graph and change the edges. However, the number of iterations that label propagation is allowed to run is gradually increased from 1, based on the ratio of the edge weights in the original graph and the edge weights of the new added edges. This is done since we expect the graph to converge towards the truth and therefore labels from the current node should ideally be propagated to all highly connected neighbours. At the end of label propagation, each label associated with a contig has a weight in the range (0, 1], and each contig may have up to a maximum of 3 labels (a limitation of the implementation of the *absorption* algorithm we adopt).

### Differential expression

To test the ability of different methods in detecting differentially expressed genes, we use contig level quantification obtained via Sailfish. The counts are aggregated to the gene-level using the R package, tximport [18], with the “scaledTPM” option. To obtain the “true” gene-level expression, the contig-to-gene aggregation is done using on the contig-to-gene mapping obtained from the genome based analysis. A differential expression analysis is then performed using `DESeq2` [19] and genes with an adjusted p-value of ≤ 0.05 are considered as differentially expressed (DE).

For the other methods analysed, the same contig level counts from Sailfish are used, to avoid bias, and the same procedure repeated using `tximport` and `DESeq2`, except that the contig-to-gene mapping is based on the clustering obtained via each method. For the FDR-sensitivity plot, the false discovery rate is controlled using the adjusted p-value cutoff for each method, and calculating the proportion of truly DE genes detected at that p-value (where all genes called DE at an FDR ≤ 0.05 under the genome based contig-to-gene mapping are considered as truly DE). For the recall-precision plot, the number of true and false positives detected at each adjusted p-value are used. Note that this plot is only for genes with adjusted p-value less than or equal to 0.05 and, therefore, the total number of genes called differentially expressed (true and false) varies for each method.

### Data Used

The first dataset used (SRA accessions SRR493366-SRR493371) is from human, *Homo sapiens*, primary lung fibroblast samples, with and without a small interfering RNA (siRNA) knock down of HOXA1 [21]. Differential expression testing was done on this dataset as well. The second sample is from dendritic cells in mice, *Mus musculus* (SRA accession SRR203276) [25]. The last dataset is from deep sequencing of Asian rice, *Oryza sativa* (SRA accessions SRR037735-SRR037738) [23]. The genomes used to obtain the “true” clustering for these experiments were hg19, GRCm38 and IRGSP-1.0 for human, mouse and Asian rice, respectively. The genomes and annotations for all the related species were obtained from the Ensembl databse [24]. Related species used for results in the main paper are macaque (*Macaca mulatta*, assembly MMUL 1) for human, rat (*Rattus norvegicus*, assembly Rnor 6.0) for mouse, and red rice (*Oryza punctata*, assembly AVCL00000000) for Asian rice. Other results in the supplementary material include those using chimp (*Pan troglodytes*, assembly CHIMP2.1.4), organutan (*Pongo abelii*, assembly PPYG2), gorilla (*Gorilla gorilla gorilla*, assembly gorGor3.1), gibbon (*Nomascus leucogenys*, assembly Nleu1.0), mouse lemur (*Microcebus murinus*, assembly micMur1), kangaroo rat (*Dipodomys ordii*, assembly dipOrd1) and wild rice (*Oryza barthii*, assembly ABRL00000000).

### Experimental Setup

All experiments listed in this paper are performed on a 64-bit linux server, running Ubuntu 14.04, with 256GB of RAM and 4 x 6-core Intel Xeon E5-4607 v2 CPUs (with hyper-threading) running at 2.60GHz. Commands used for the experiments are explained in the supplementary material.

